# Late-Stage Skeletal Muscle Transcriptome in Duchenne muscular dystrophy shows a BMP4-Induced Molecular Signature

**DOI:** 10.1101/2024.04.19.590266

**Authors:** Hanna Sothers, Xianzhen Hu, David K. Crossman, Ying Si, Matthew S. Alexander, Merry-Lynn N. McDonald, Peter H. King, Michael A. Lopez

**Author notes:** Corresponding Author: Michael A. Lopez, M.D., Ph.D., Address: Department of Pediatrics, University of Alabama at Birmingham, CHB314, 1600 7th Avenue South, Birmingham, AL 35233 USA., Peter H. King, M.D., Address: Department of Neurology, University of Alabama at Birmingham, Civitan 545C, 1530 3rd Avenue South, Birmingham, AL 35294 USA.

## Abstract

Duchenne muscular dystrophy (DMD) is a fatal X-linked recessive disease due to loss-of-function mutations in the *DYSTROPHIN* gene. DMD-related skeletal muscle wasting is typified by an aberrant immune response involving upregulation of TGFβ family of cytokines. We previously demonstrated that bone morphogenetic protein 4 (BMP4) is increased in DMD and BMP4 stimulation induces a 20-fold upregulation of *Smad8* transcription. However, the role of BMP4 in severely affected DMD skeletal muscle is unknown. We hypothesized that transcriptomic signatures in severely affected human DMD skeletal muscle are driven by BMP4 signaling. Transcriptomes from skeletal muscle biopsies of late-stage DMD vs. non-DMD controls and C2C12 muscle cells with or without BMP4 stimulation were generated by RNA-Seq and analyzed for single transcript differential expression as well as by Ingenuity Pathway Analysis and weighted gene co-expression network analyses. A total of 2,328 and 5,291 transcripts in the human muscle and C2C12 muscle cells, respectively, were differentially expressed. We identified an overlapping molecular signature of 1,027 genes dysregulated in DMD muscle that were induced in BMP4-stimulated C2C12 muscle cells. Highly upregulated DMD transcripts that overlapped with BMP4-stimulated C2C12 muscle cells included *ADAMTS3, HCAR2, SERPING1, SMAD8*, and *UNC13C.* The DMD transcriptome was characterized by dysregulation of pathways involving immune function, extracellular matrix remodeling, and metabolic/mitochondrial function. In summary, we define a late-stage DMD skeletal muscle transcriptome that substantially overlaps with the BMP4-induced molecular signature in C2C12 muscle cells. This supports BMP4 as a disease-driving regulator of transcriptomic changes in late-stage DMD skeletal muscle and expands our understanding of the evolution of dystrophic signaling pathways and their associated gene networks that could be explored for therapeutic development.

## Introduction

Duchenne muscular dystrophy (DMD) is a fatal X-linked recessive disease caused by loss of dystrophin expression^1^. This triggers a cascade of events starting with destabilization of the sarcolemmal membrane of skeletal muscle. Consequently, a chronic immune response ensues, characterized by upregulation of the transforming growth factor (TGF) receptor family signaling^2^. As the disease progresses to late-stage, skeletal muscle undergoes fibroadipocytic conversion and ends in total paralysis with loss of ambulation, respiratory failure, and death in the third to fourth decades. Anti-inflammatory therapies have been helpful but unsatisfactory due to side effects and modest impact on progression and survival^3^. The large size of *DYSTROPHIN* (2.5 Mb) is a major barrier to restoring dystrophin expression, and as such, dystrophin restoration therapies have had limited success^1,4^. Therefore, identification of disease-driving signaling pathways remains an important area for expanding treatment of DMD^5^.

TGFβ signaling via canonical TGFβ receptor ligands is well-recognized, but the role of BMP mediated pathology is underexplored^5–7^. We have shown that BMP4 is highly upregulated in severely affected DMD skeletal muscle in parallel with *SMAD8* (officially named *SMAD9*), a TGF intracellular transcription factor^8^. We further showed that SMAD8 is strongly activated by BMP4 in C2C12 muscle cells and exerts a negative effect via SMAD8 on myogenesis associated with a suppression of muscle-enriched microRNAs (myomiRs) miR-1, miR-133a, and miR-133b^8^. MyomiRs are broad regulators of genes expression and extend the regulatory reach of SMAD8. Investigation of disease-driving pathways, regulators, and networks, including BMP4, in severely affected late-stage DMD muscle has been slowed by decreasing use of muscle biopsy and difficulty with animal models reproducing severe dystrophic muscle disease seen in humans. As such, prior studies have largely consisted of array panels in younger DMD muscles^9–14^.

To begin addressing these challenges, we generated RNA-Seq transcriptomic data followed by Ingenuity Pathways Analysis (IPA) and Weighted Gene Correlation Network Analysis (WGCNA) to characterize the late-stage human DMD skeletal muscle transcriptome. We compared findings to previously reported gene profiling studies to identify stage-specific patterns. We then compared late-stage DMD muscle transcriptome to that of BMP4-stimulated C2C12 muscle cells and found a substantial overlap, thus revealing a novel role of BMP4 signaling in shaping the transcriptome in late-stage DMD.

## Methods

### Human Muscle Samples

Human muscle biopsies were collected from 3 DMD patients under an approved UAB Institutional Review Board protocol (IRB00000196). 3 human non-DMD muscle samples were selected from an archive of remnant muscle biopsy tissues in the UAB Division of Neuromuscular Disease without dystrophic pathology as previously detailed^15^. The non-DMD ranged in age from 8 to 59 years old and DMD male patients ranged in age from 15 to 20 years old. Non-DMD samples (2 male, 1 female) were taken from the deltoid, tibialis anterior, and quadriceps muscles. The DMD muscle samples were taken from paraspinous muscle from non-ambulatory males undergoing scoliosis surgery. DMD patients all had out-of-frame mutations with Duchenne phenotype. 1 patient was on daily prednisone and 2 were steroid naive.

### Cell culture

BMP4 stimulation was performed as previously published^8^. Briefly, C2C12 muscle cells were grown in growth media on 6-well plates seeded with 3 x 10^5^ cells per well 24 h prior to stimulation. BMP4 recombinant protein (R&D System/Thermo Fisher Scientific, Waltham, MA, USA; Ref# 5020-BP) was used at 400 ng/mL. C2C12 cells were originally obtained from ATCC (Cat# CRL-1772).

### Sequencing and RNA-Seq analysis

DNALink performed skeletal muscle total RNA isolation using the Truseq Stranded Total RNA H/M/R Preparation Kit and Next Generation Sequencing using the Illumina NovaSeq6000 platform. Pre-processing was performed using FastQC. STAR (version 2.710a) was used to align reads to the reference genome (Gencode GRCm39 Release M26) (using parameters - outReadsUnmapped Fastx-outSAMtype BAM SortedByCoordinate-outSAMattributes ALL)^16^. Transcript abundances were calculated using HTSeq-count (version 2.0.2; using parameters -m union -r pos -t exon -i gene_id -a 10 -s no -f bam)^17^. Transcripts were aligned with all 6 samples having > 85% uniquely mapped alignments. DESeq2 was used to calculate differential gene expression (**Table S0A, Table S0B**)^18^. Log_2_ Fold Change (LFC) and standard error show gene’s expression changed due to disease state. The basic annotation performed used gene annotation from NCBI db. An outlier analysis for replicated “BMP4-3” showed no skewing of P-Values or log fold change (**Fig. 2S**).

### Ingenuity Pathway Analysis

Canonical Pathways were generated through use of QIAGEN Ingenuity Pathway Analysis (Qiagen, Redwood City, CA) as previously described^19–21^. CP analysis identified the pathways from the QIAGEN Ingenuity Pathway Analysis library of canonical pathways that were most significant to the data set. CP’s likelihood of activation or inhibition for each pathway is calculated as a Z-score which is based on enrichment of genes overlapping with Ingenuity Knowledge Database CPs activated or inhibited state. P-Value is based on enrichment of genes overlapping with Ingenuity Knowledge Database. Molecules from the data set that met FDR < 0.05 and fold change > |2.0| and were associated with a CP in the QIAGEN Knowledge Base were considered for the analysis. The significance of the association between the data set and the CP was measured in two ways: 1) A ratio of the number of molecules from the data set that map to the pathway divided by the total number of molecules that map to the CP is displayed; and 2) A right-tailed Fisher’s Exact Test was used to calculate a P-Value determining the probability that the association between the genes in the dataset and the CP is explained by chance alone. 3) In many cases a Z-score was calculated to indicate the likelihood of activation or inhibition of that pathway.

### RNA Isolation and qPCR Analysis

Total RNA was isolated from muscle tissue as previously described^8^. cDNA was synthesized using SuperScript™ IV First-Strand Synthesis System (Invitrogen/Thermo Fisher Scientific, Waltham, MA, USA) using 1 ug of RNA. Taqman assay probes and primers (Applied Biosystems/Thermo Fisher Scientific, Waltham, MA, USA). TaqMan reactions were performed using TaqMan™ Gene Expression Master Mix (Applied Biosystems/Thermo Fisher Scientific, Waltham, MA, USA). Samples were run on a ViiA 7 Real-Time PCR System (Applied Biosystems/Thermo Fisher Scientific, Waltham, MA, USA) in 386 well plates. Relative expression values were calculated using the ΔΔCT method with normalization to the housekeeping gene.

### Transcriptomics analyses

Weighted Gene Correlation Network Analysis (WGCNA) is a well-established tool for creating networks of co-expressed biomarkers, such as RNA transcripts^22,23^. Instead of using established biological pathways, the networks are generated using an agnostic approach, allowing for a wider range of transcripts to be identified. It can also be used to test for preservation of co-expression patterns between independent transcriptomics datasets, enabling validation of key players in disease processes.

The WGCNA R package was used to identify co-expressed RNA transcripts and assess module preservation of the DMD network within the C2C12 network. The DMD data was filtered for low counts (n<10 reads across all samples) prior to normalization with DESeq2^18^. The variance stabilized expression data was used for WGCNA. Signed correlation networks were built using a Bi-weight Mid-correlation. Soft thresholding powers (β) of 10 for the DMD network and 16 for the C2C12 network were calculated using approximate scale-free topology. WGCNA uses hierarchical clustering to separate the transcripts into modules. Modules with ≥70% similarity were merged. The average expression profile of a module is represented by its eigengene. The eigengenes are used to assess the correlation of modules to a particular trait (disease status, age, sex, muscle group). Modules with a P-Value <0.05 were significantly correlated to a trait. In addition to this, the correlation of a gene’s expression to a module’s eigengene is known as kME and is used a measure of module membership. This value is also highly correlated with intra-modular connectivity. So, it’s also used to evaluate the “hubness” of a transcript ^73^. In networks, hub nodes are connected to many other nodes. A cutoff of kME >0.8 was used to identify hub genes. The top hub genes for each module were selected using WGCNA’s built in function “chooseTopHubInEachModule”.

Preservation of human genes in the C2C12 modules was carried out on a subset of transcripts with a one-to-one mapping between human and mouse Ensembl IDs^24^. Module preservation calculates different Z-statistics that are summarized into the Z-Summary statistic. A value greater than 2 indicates moderate preservation and a value greater than 10 indicates high preservation.

### Network Analysis

Files containing the node and edge data from the resulting topological overlap adjacency matrices were generated using WGCNA where an edge weight represents correlation strength between two transcripts and node size corresponds to the degree of connectivity (hubness) of an individual transcript to other transcripts in the network. Edges below a set cutoff weight were excluded. Cutoffs were selected based on the distribution of edge weights in the module. Cutoffs for individual modules were selected based on the edge weight distribution as follows: Turquoise - 0.400 (human) and 0.441 (C2C12), Blue - 0.450 (human) and 0.387 (C2C12). Cytoscape (version 3.10.0) was used to visualize the networks. In a network, first neighbors are nodes that share an edge. First neighbor networks were generated for relevant genes from the adjacency matrices exported from WCGNA. Genes with many connections were limited to the top 10 - 20 nodes with the highest edge weight.

## Results

### Differentially Expressed Transcripts in Human DMD late-stage skeletal muscle

RNA-Seq data was generated and analyzed in 3 DMD and 3 non-DMD patients. Principal component (PC) analysis demonstrated stratification of the DMD from the non-DMD muscles using the first two PCs (**Fig. 1SA**).

A total of 2,328 transcripts were differentially expressed at a false discovery rate (FDR) < 0.05 (**Table S1**). Among the protein encoding transcripts, 374 were upregulated and 172 were downregulated with log_2_ fold-change (LFC_DMD_) ≥ |2.0| (**Fig. 1A**). The top transcript upregulated in DMD muscle was *HCAR2* [LFC_DMD_ = 6.29 ± 1.44, FDR = 5.19 x 10^-04^], which encodes the Hydroxycarboxylic acid receptor 2, a member of the G-protein-coupled receptor family^25^. *HLA-A*, major histocompatibility complex (MHC), class I, A, was one of the most downregulated [LFC_DMD_ = −7.07 ± 0.86, FDR = 5.34 x 10^-13^]. Other TGF and immune-related factors included upregulation of *TGFB1* upregulated [LFC_DMD_ = 1.96 ± 0.423, FDR = 2.14 x 10^-04^ and *SMAD8* was the only R-Smad upregulated [LFC_DMD_ = 1.93 ± 0.626, FDR = 2.09 x 10^-02^]. Additionally, key regulatory factors in myogenesis were altered including downregulation of *MEF2D* [LFC_DMD_ = −1.09 ± 0.355, FDR = 2.15 x 10^-02^], downregulation of *MYF6* [LFC_DMD_ = −1.4 ± 0.327, FDR = 7.56 x 10^-04^], and upregulation of *MYOG* [LFC_DMD_ = 1.28 ± 0.256, FDR = 5.11 x 10^-05^]. *UNC13C*, involved in neuromuscular junction function, was downregulated [LFC_DMD_ = −6.39 ± 2.25, FDR = 3.67 x 10^-02^]. Markers of muscle regeneration, embryonic and fetal myosin heavy chain transcripts (*MYH3*, *MYH8*) were also upregulated. Further, several tissue and extracellular matrix (ECM) remodeling genes were dysregulated. A Disintegrin and Metalloprotease Domain (ADAM) metalloproteinases were broadly upregulated including *ADAM12*, 2*2, 28 and ADAM* thrombospondins (*ADAMTS*) *2*, 3, *7, 12, 15*, and *16*. Similarly, matrix metalloproteinases (MMP) were broadly upregulated (*MMP2*, *9*, *14*, *16*, *19*) with *MMP9* being the highest [LFC_DMD_ = 3.1 ± 1.03, FDR = 2.45 x 10^-02^]. *DMD* [LFC_DMD_ = −2.0 ± 0.29, FDR = 3.69 x 10^-09^] and *CKM* [LFC_DMD_ = −2.83 ±, FDR = 7.45 x 10^-05^] transcripts were significantly downregulated.

**Fig. 1.**
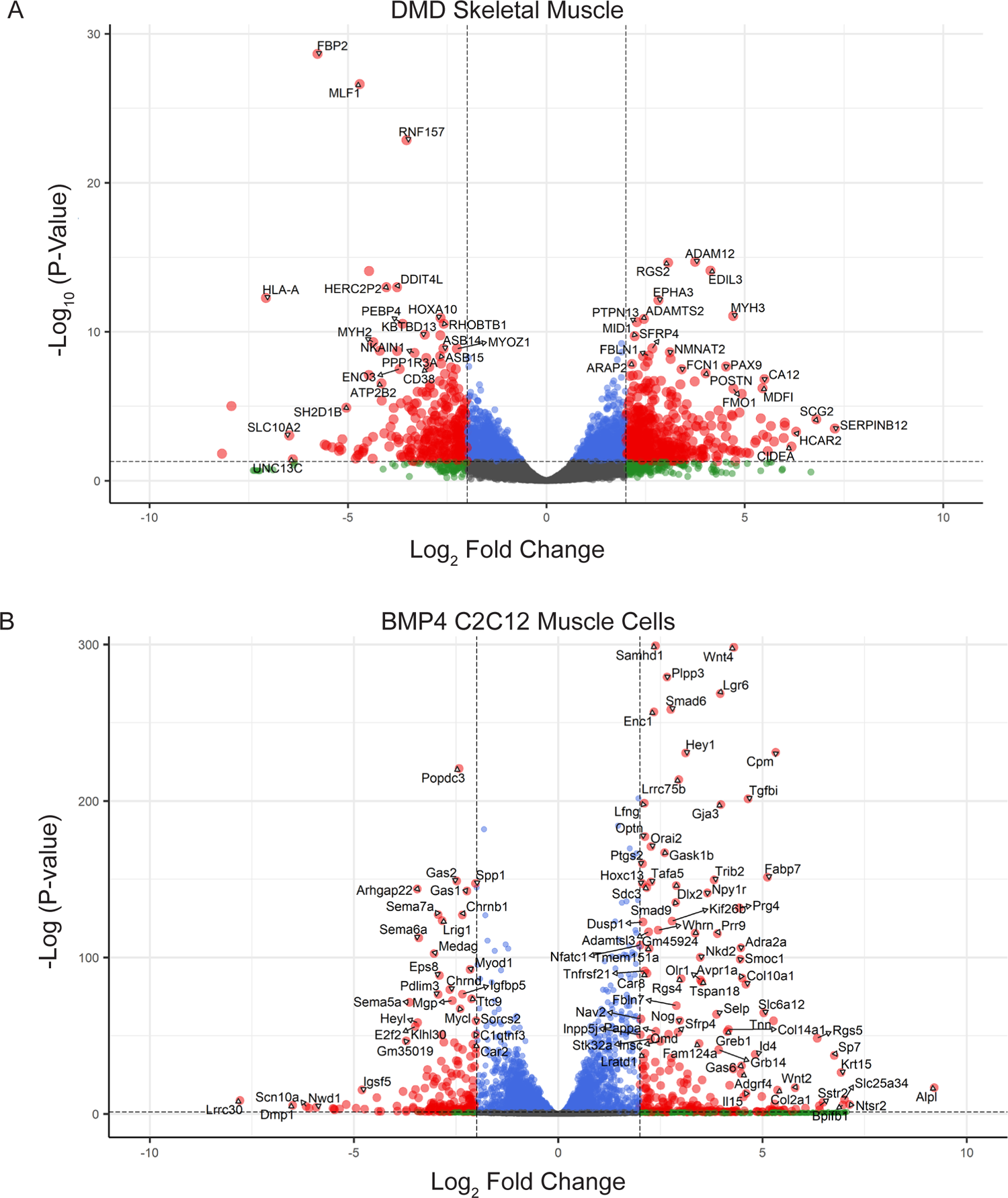
RNA-seq analysis showing differential gene expression of DMD muscle and BMP4-stimulated C2C12 muscle cells. (A) Volcano plot of DMD skeletal muscle transcriptome showing -log_10_ of adjusted P-Value vs. log_2_ fold change. Dashed vertical lines mark log_2_-fold change > |2|. Dashed horizontal line marks adjusted P-Value < 0.05. Red indicates significant and > |2| log_2_-fold change. Blue indicates significant but log_2_-fold change < |2|. Green indicates not significant and log_2_-fold change > |2|. Grey indicates not significant and log_2_-fold change < |2|. (B) Volcano plot of BMP4-stimulated muscle cells (C2C12) transcriptome showing -log_10_ of adjusted P-Value vs. log_2_ fold change. Dashed vertical lines mark log_2_-fold change > |2|. Dashed horizontal line marks adjusted P-Value < 0.05. Red indicates significant and > |2| log_2_-fold change. Blue indicates significant but log_2_-fold change < |2|. Green indicates not significant and log_2_-fold change > |2|. Grey indicates not significant and log_2_-fold change < |2|.

### BMP4-stimulated muscle cell transcriptome

We then analyzed BMP4-stimulated C2C12 muscle cells using RNA-Seq. Principal component analysis demonstrated stratification of the BMP4-stimulated muscle cells (**Fig. 1SB**). In the BMP4-stimulated transcriptome 5,061 transcripts were differentially expressed, compared to vehicle control, with an FDR < 0.05 (**Table S11**). Of these, 202 upregulated and 139 downregulated transcripts had LFC_BMP4_ ≥ |2.0| (**Fig. 1B**).

Alkaline phosphatase, *Alpl*, was the highest upregulated transcript [LFC_BMP4_ = 9.2 ± 1.04, FDR = 1.4 x 10^-17^]. Neurotensin receptor 2, *Ntsr2*, was the second highest upregulated transcript [LFC_BMP4_ = 7.1 ± 1.34, FDR = 9.6 x 10^-07^]. It functions in apoptosis and cell viability. The most significantly downregulated genes included *Lrrc30* [LFC_BMP4_ = −7.8 ± 1.22, FDR = 2.41 x 10^-09^]. Its function is unknown, but speculated to have similar actin and alpha-actinin-binding based on its high expression in skeletal muscle (dbGaP Accession: phs000424.v8.p2) and structural homology to *LRRC10^26^*. There were several genes involved in proliferation and cell cycle control that were downregulated including *Ckd6*, the most downregulated cyclin dependent kinases, [LFC_BMP4_ = −1.5 ± 0.14, FDR = 3.78 x 10^-28^]. *Mki67* [LFC_BMP4_ = −0.90 ± 0.11, FDR = 2.1 x 10^-14^] and *Top2a* [LFC_BMP4_ = −1.0 ± 0.08, FDR = 2.3 x 10^-38^] were highly downregulated proliferation markers. *E2f2* [LFC_BMP4_ = −3.5 ± 0.22, FDR = 1.0 x 10^-56^] was the most highly downregulated E2 transcription factor.

### Ingenuity Pathway Analysis (IPA) of DMD Transcriptomes

#### Top canonical pathways and regulators dysregulated in DMD muscle

The differentially expressed transcripts in the DMD case-control analysis (FDR < 0.05) encompassed several canonical pathways (CPs) (**Fig. 2A, Table S2**). Activated CPs included Mitochondrial Dysfunction, Role of Osteoclasts in Rheumatoid Arthritis Signaling, Neutrophil Degranulation, Phagosome Formation, Integrin cell surface interactions, and Extracellular Matrix Organization. For example, the Mitochondrial Dysfunction CP [Z-score = 4.7, P-Value = 4.1 x 10^-24^] was predicted to be highly activated with downregulation of numerous Complex I, II, III, IV, and V genes which was also reflected in inhibited pathways like the Electron Transport Chain, Oxidative Phosphorylation, The Citric Acid Cycle, and Mitochondrial Translation.

**Fig. 2.**
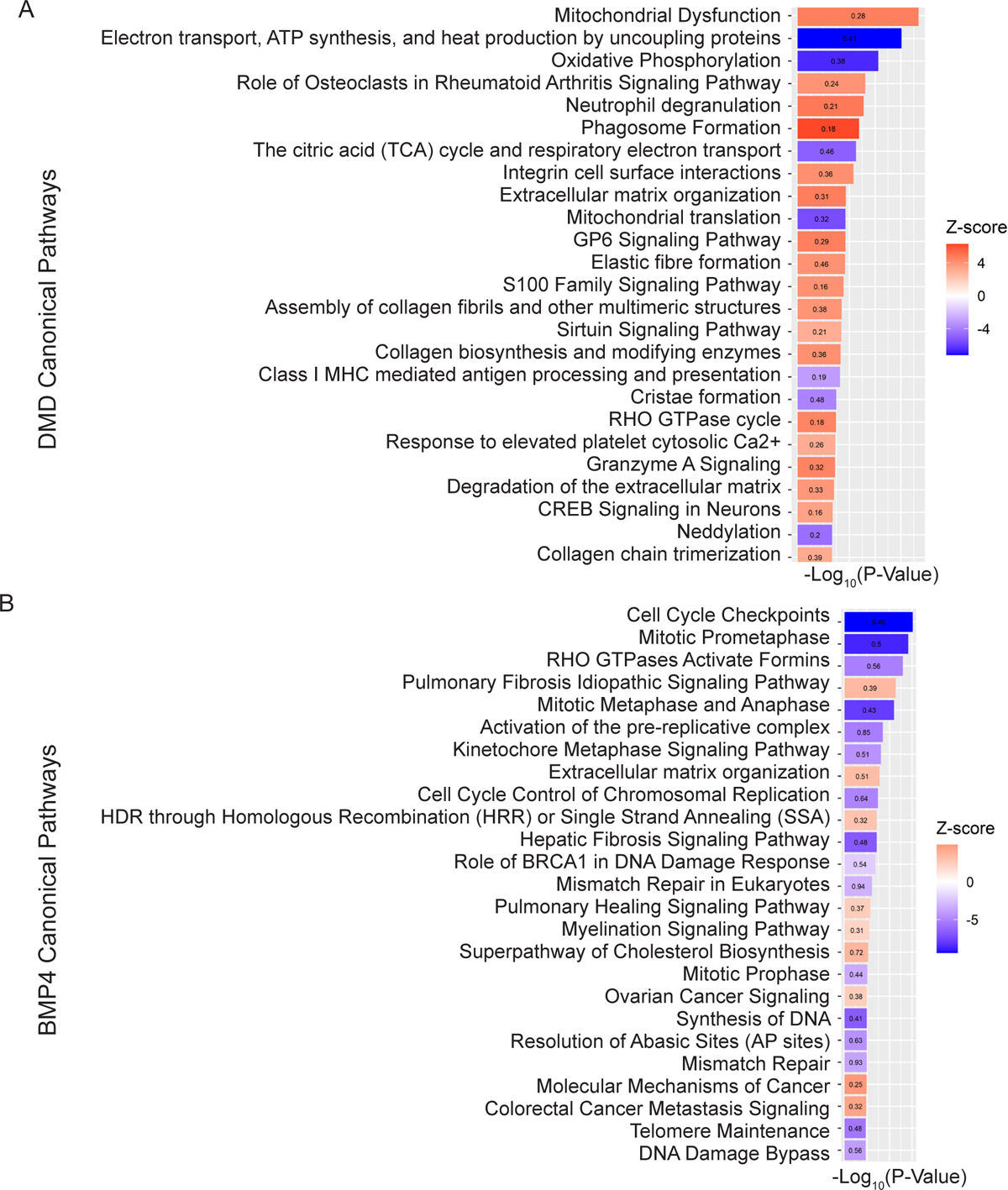
Ingenuity Pathway Analysis (IPA) predicted activated or inhibited Canonical Pathways. (A) DMD skeletal muscle IPA canonical pathways. Heat maps show Z-scores for each canonical pathway ordered by -log_10_ of adjusted P-Value. The top 25 canonical pathways with Z-scores > |3.0| were selected for comparison. Bars are labelled with the ratio of the number of overlapping genes within the canonical pathway. (B) BMP4-stimulated C2C12 muscle cell canonical pathways predicted by Ingenuity Pathway Analysis (IPA). Heat maps show Z-scores for each canonical pathway ordered by -log_10_ of adjusted P-Value. The top 25 canonical pathways with Z-scores > |2| were selected for comparison. Bars are labelled with the ratio of the number of overlapping genes within the canonical pathway. Z-scores statistics show predicted state as activated (red) or inhibited (blue).

Ingenuity Pathway Analysis (IPA) also identified upstream regulators that were predicted to be activated or inhibited based on gene expression changes of transcriptional target molecules in each transcriptome (**Table S3)**. There were 84 upstream regulators with predicted Z-scores of more than |3.5|. TP53, TGFβ1, TNF, and DMD were amongst the top 30 by P-Value (**Fig. 3A**).

**Fig. 3.**
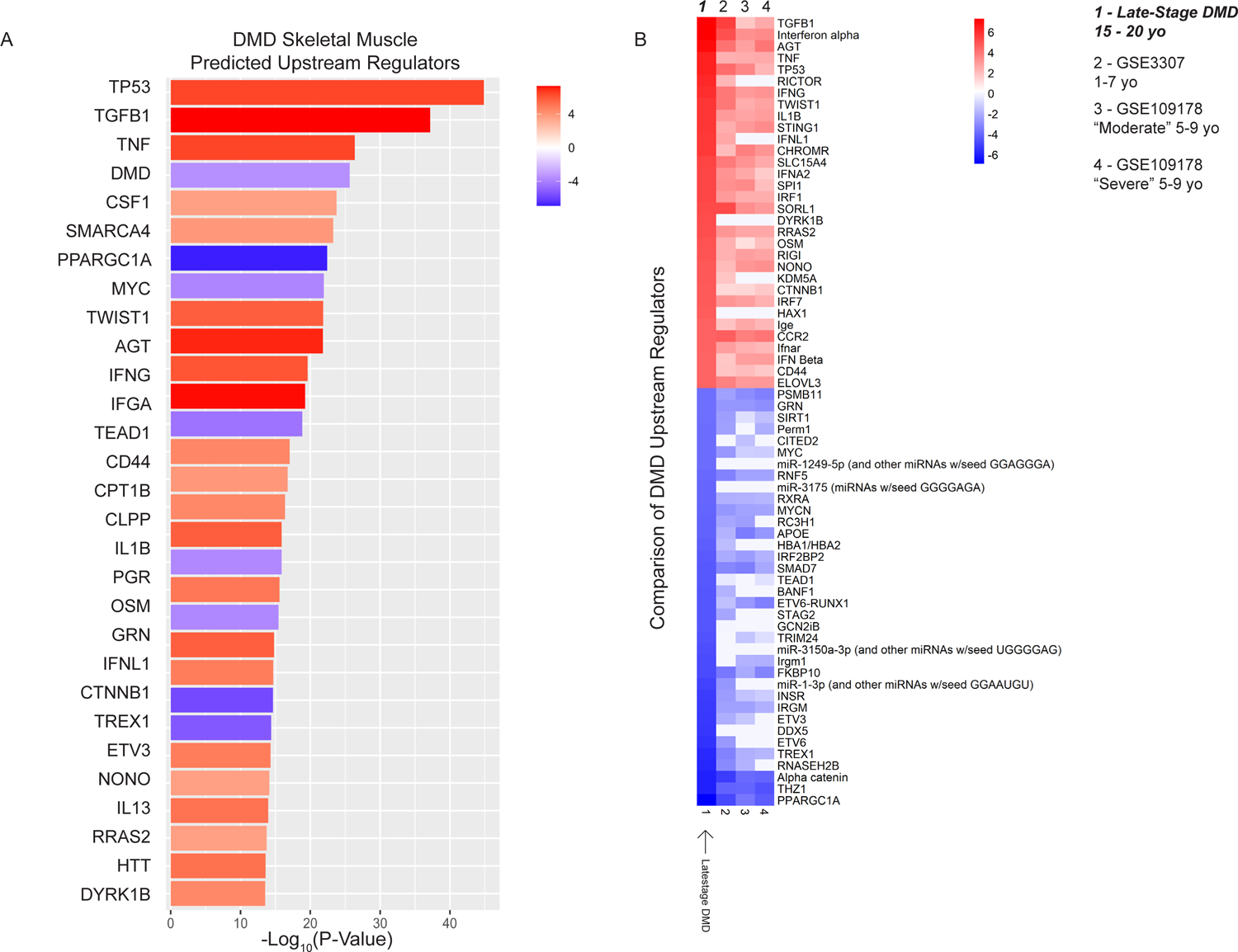
DMD skeletal muscle Ingenuity Pathway Analysis (IPA) predicted upstream regulators. (A) DMD upstream regulators are shown in heat maps with Z-scores for each upstream regulator ordered by -log_10_ of adjusted P-Value. The top 30 upstream regulators with Z-scores > |3.5| were selected for comparison. Z-scores statistics show predicted activated (red) or inhibited (blue) upstream regulators. (B) Heat map shows comparison of predicted upstream regulators from late-stage DMD skeletal muscle transcriptome (1) with younger DMD muscle arrays that were identified by analysis match (2) - GSE3307, (3) - GSE109178 / Moderate, (4) - GSE109178 / Severe). Z-scores statistics show predicted state as activated (red) or inhibited (blue).

#### Comparison of late-stage DMD transcriptome to younger IPA-analysis matched databases

Using analysis match, previously published DMD array-based data (GSE3307, GSE109178) showed similar upstream regulators predicted to be dysregulated. Our DMD data (DMD Non-Amb) matched to datasets of dystrophic muscle from younger DMD patients (**Fig. 3B, Table S4**)^14,27^. Several cytokines were predicted to be activated upstream regulators including TGFβ1, TNF, IFNα/γ, and IL1B. In our older DMD muscles, TGFβ1 had higher predicted Z-score activations (7.3) compared with GSE3307 (5.5) and GSE109178 (1.8, 2.5). TP53, IFNα/γ, TNF and IL1B showed reversal of inhibition to an activated state in more severely affected late-stage DMD tissue. Common upstream regulators predicted to be inhibited included SMAD7 and RNF5. PPARGC1A, alpha catenin, CITED2, and miR-1-3p showed increased inhibition in the late-stage DMD muscles in contrast to younger DMD samples. *DYRK1B*, *HAX1*, miR-1249-5p, and *DDX5* were newly predicted activated upstream regulators in late-stage DMD muscle.

### Ingenuity Pathway Analysis (IPA) of BMP4-Stimulated Muscle Cell Transcriptomes

#### Canonical Pathways in BMP4-Stimulated Muscle Cells

In the BMP4-stimulated muscle cells, there were several CPs predicted to be activated or inhibited by BMP4 stimulation (**Fig. 2B, Table S12**). The activated CPs included Pulmonary Fibrosis Idiopathic Signaling Pathway, ECM Organization, and HDR through Homologous Recombination or Single Stran Annealing. The inhibited CPs included Cell Cycle Checkpoint (CCC), Mitotic Prometaphase, RHO GTPases Activate Formins, and Mitotic Metaphase and Anaphase. The CCC, for example, showed downregulation of numerous genes such as cyclin dependent kinases, cyclin dependent kinase inhibitors, and tumor suppressor genes (*CDK1*, *CKD2*, *CDKN1A*, and *TP53*). The CCC pathway is involved in tight control of DNA replication. Early differentiating myoblasts show inhibition of this pathway compared with late differentiating myoblasts. In the Pulmonary Fibrosis CP, CTNNB1, WNT1, IL1B, and STAT6 were predicted activated. These are involved in ECM accumulation and fibroblast proliferation.

#### Common DMD and BMP4 Pathways

Comparing the DMD and BMP4 transcriptomes we found common dysregulated CPs (**Table S2, Table S12**). The shared CPs included Phagosome Formation, Molecular Mechanisms of Cancer, and S100 Family Signaling Pathway. The Phagosome Formation CP, for example, showed higher Z-scores [DMD Z-score = 3.0, BMP4 Z-score = 6.3] indicating higher activation in both transcriptomes. There was also common activation of PIP3, AKT, MTOR, and p70 involved in phagocytosis and SYK, PPKCD, CARD9, and BCL10 involved in inflammatory response of cells. Molecular Mechanisms of Cancer CP included matrix metalloproteinases which were broadly upregulated. For example, in the BMP4-stimulated muscle cells *MMP2*, *11*, *14*, *19*, *23B*, and *28* were upregulated. While in DMD transcriptome, *MMP2*, *9*, *14*, *16*, and *19* were upregulated. Commonly downregulated pathways included Cell Cycle Checkpoints, Electron transport, ATP synthesis and heat production, Mitotic Metaphase and Anaphase, and Regulation of TP53 Activity through Phosphorylation.

#### BMP4-Stimulated Muscle Cells Predicted Upstream Regulators

In BMP4-stimulated muscle cells, the top three predicted activated protein-encoding regulators were *TGF*β*1*, *XBP1*, and *TP53* (**Table S13**). The top predicted activated regulators amongst growth factors were Bmp2, Bmp4, Bmp10, Gdf2, Tgfβ1, Tgfβ2, and Tgfβ3. The only predicted inhibited growth factors were Areg, Nog, Hgf, and Angptl3. Amongst cytokines, Il1b, Il6, Prl, and Wnt3a had the highest Z scores. While *Csf2*, *Ifnb1*, and *Il10* were the only 3 inhibited transcripts. Shared upstream regulators that were predicted to be activated or inhibited in both DMD muscle and BMP4-stimulated transcriptomes included activation of TGFβ1, TP53, and RICTOR and inhibition of MYC, CTNNA1/2, and MYCN (**Table S3**, **Table S13**).

#### Weighted gene co-expression network analysis (WGCNA) of DMD muscle transcriptome

We next used WGCNA to identify modules of co-expressed genes correlated with the DMD disease state. WGCNA identified a total of 29 modules of co-expressed genes (**Table S5**). Of these, 4 modules were significantly associated with the DMD disease state (**Fig. 4A**); turquoise (Turquoise_DMD_), blue (Blue_DMD_), cyan (Cyan_DMD_) and dark green (Dark green_DMD_). The Turquoise_DMD_ module was comprised of 5,730 genes collectively upregulated among DMD muscles with *SERPING1* as the top hub gene with a kME of 0.996 (**Table S6**). Top IPA upstream regulators in the Turquoise_DMD_ module included TGFβ1, IFNG, MMP9, IL1B, and TP53. S*MAD8* was also a member of the Turquoise_DMD_ module. The Blue_DMD_ (4,261 genes), Cyan_DMD_ (551 genes), and Dark green_DMD_ (401 genes) modules were downregulated in DMD muscle (T**able S7**). The Blue_DMD_ module, of which the DMD transcript (ENSG00000198947) is a member, was significantly downregulated (**Table S7**) and its top hub gene was *AKTIP*. Top IPA upstream regulators in Blue_DMD_ module included PPARGC1A, BANF1, and DMD. *BMP4* was a member of the salmon module which was not significantly associated with DMD disease state.

**Fig. 4.**
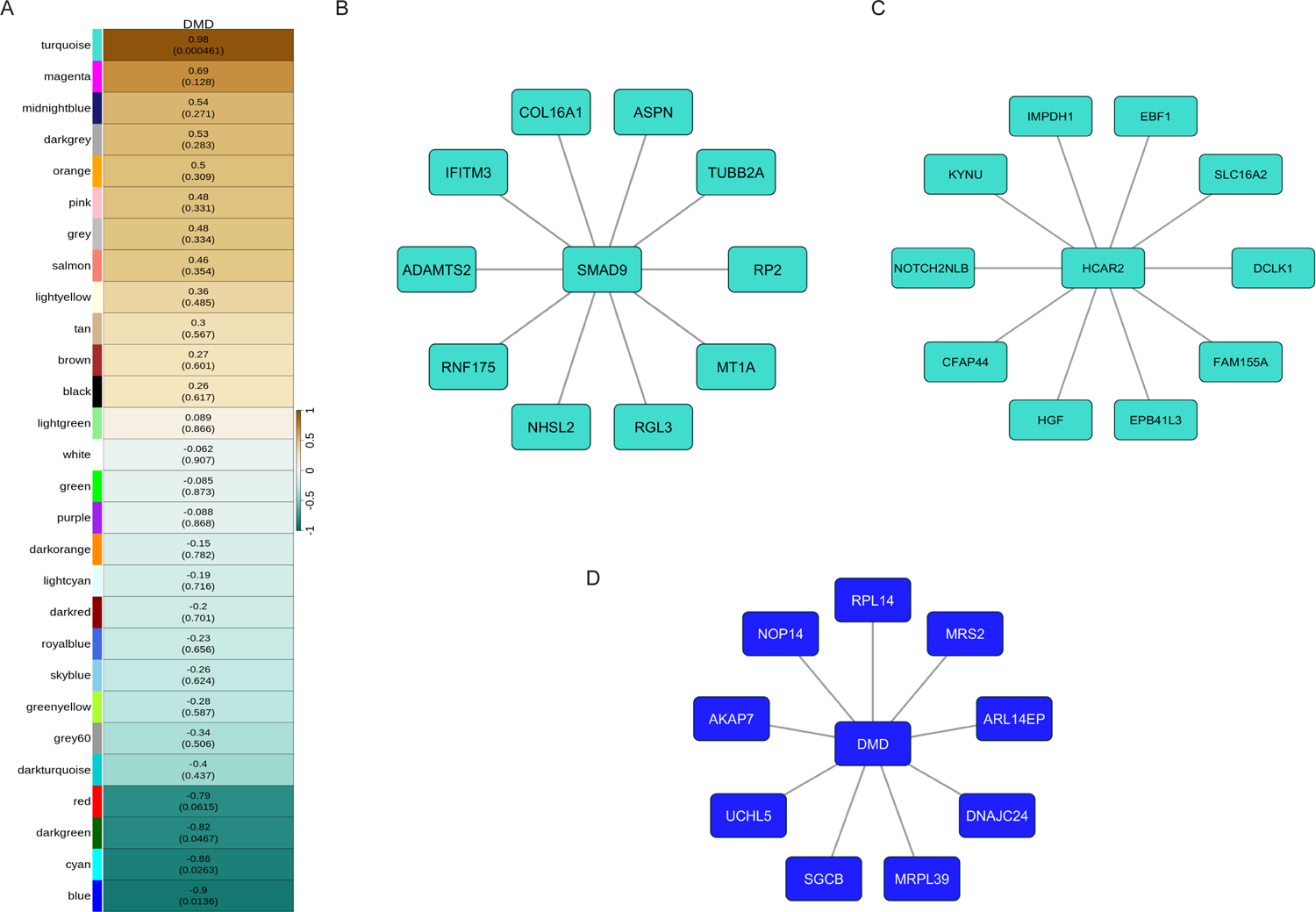
Weighted Gene Correlation Network Analysis of DMD muscle transcriptome. (A) Heat map of correlations between the module eigengenes and the DMD disease trait. Modules are labeled with Z-score and (P-Value). Z-scores indicate upregulated (+) or downregulated (-) co-expression for each module. Colors are arbitrarily assigned. (B) Top connected genes co-expressed with *SMAD8* with edge weight (from adjacency matrix) ≥ 0.40 in the Turquoise_DMD_ (upregulated) module. (C) *HCAR2* co-expressed genes with edge weights (from adjacency matrix) ≥ 0.40. (D) Top 10 genes co-expressed with the DMD transcript in downregulated Blue_DMD_ module based on edge weight *≥* 0.45.

#### Top SMAD8 Connected Genes

Deeper investigation into the genes with the highest co-expression to *SMAD8* in the Turquoise_DMD_ module revealed 16 genes with an edge weight higher than the cutoff (**Fig. 4B, Table S8**). *COL16A1* had the highest edge weight and therefore the strongest correlation to *SMAD8.* The COL16A1 gene encodes Collagen Type XVI α1 Chain which contributes to tissue structure and strength in the extracellular matrix, particularly in connective tissues involved in fibrosis and wound healing.

#### Top HCAR2 Connected Genes in Turquoise Module

We next investigated genes highly connected to *HCAR2*, the highest overall upregulated DMD transcript. A total of 46 genes had edge weights above the cutoff. *HGF, EBF1, SLC16A2, EPB41L3, FAM155A, KYNU,* and *CFAP*44 were the top genes with the highest edge weight for *HCAR2 (***Fig. 4C, Table S9**).

#### Top DMD Connected Genes

Edge weights generated from the adjacency matrix were used to identify highly connected genes to DMD within the downregulated Blue_DMD_ module (**Table S10)**. The top 10 first neighbor genes are shown (**Fig. 4D**). Mitochondrial ribosomal protein L39 (*MRPL39*) was the highest kME module membership score. While *NOP14*, is a nucleolar protein involved in ribosomal RNA processing, had the strongest edge weight score to DMD. *SGCB* is a constituent of the Dystrophin Associated Protein Complex (DAPC), and it had the largest LFC down. *UCHL5* is a deubiquitinating enzyme involved in halting protein degradation and regulates TGFβ signaling via physical interaction with SMAD2, SMAD3, and SMAD7^28^.

#### Weighted gene co-expression network analysis (WGCNA) of BMP4-stimulated myoblast transcriptome

WGCNA identified 25 modules of co-expressed genes in the C2C12 muscle cell transcriptome (**Table S14**). The Turquoise_BMP4_ (4,651 transcripts upregulated) and Blue_BMP4_ (4,452 transcripts downregulated) modules were identified as significantly correlated with BMP4 stimulation (**Fig. 5A, Table S15-S16**). *Angptl2* and *Rfc3* and were the associated hub genes for Turquoise_BMP4_ and Blue_BMP4_ modules, respectively.

**Fig. 5.**
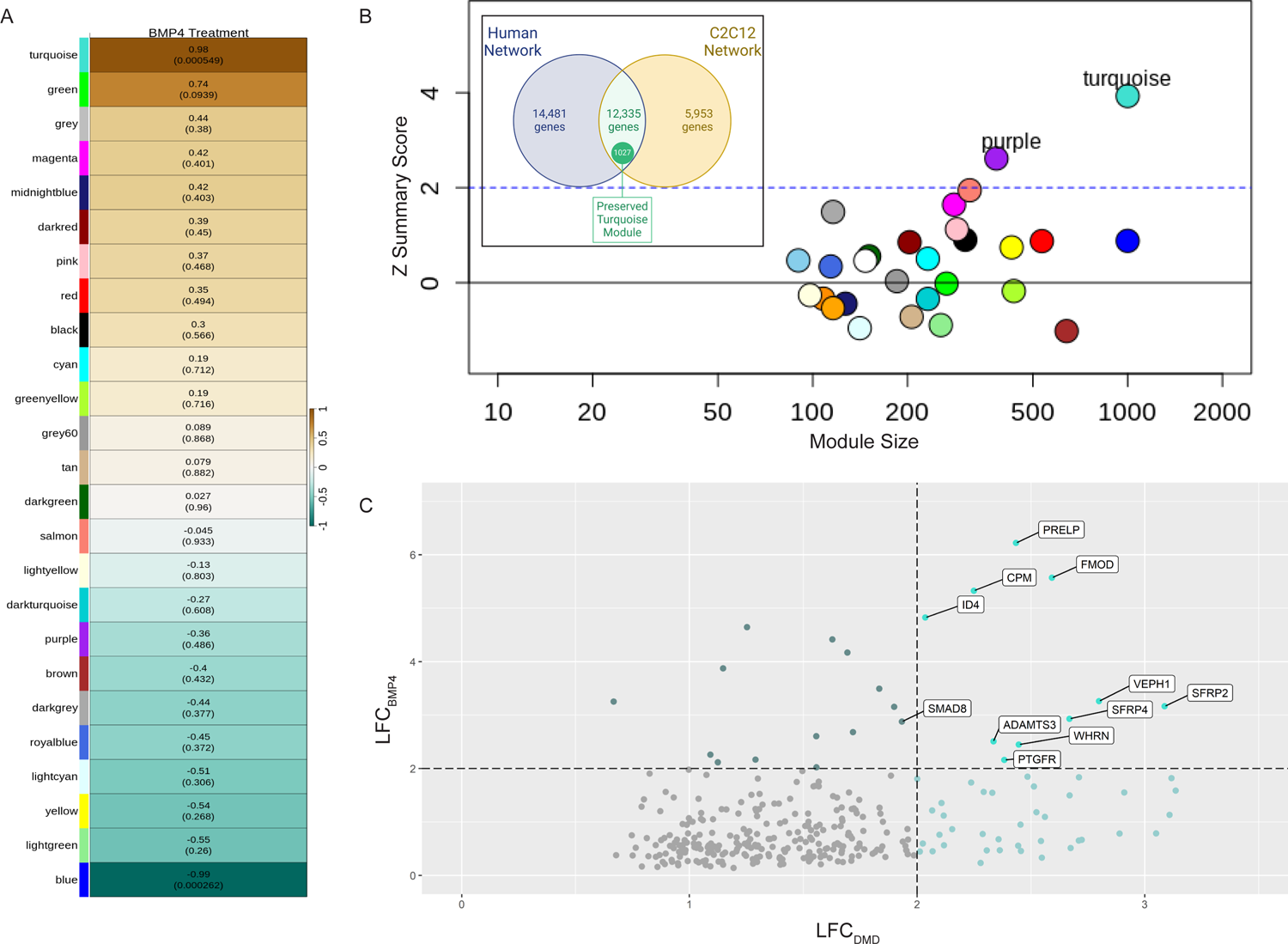
Weighted Gene Correlation Network Analysis of BMP4-stimulated C2C12 transcriptome and comparison to DMD transcriptome. (A) Heat map showing correlation between the module eigengenes and BMP4-stimulated muscle cells. Modules are labeled with Z-score and (P-Value). Z-scores indicate upregulated (+) or downregulated (-) co-expression for each module. Colors are arbitrarily assigned. (B) Module preservation analysis between the DMD and BMP4-stimulated C2C12 muscle cells showing Z-Summary preservation statistic for each module and module size. The Turquoise_preserved_ module was moderately preserved with a Z-Summary score of 4.7. Inset shows Venn diagram of overlapping genes in the Turquoise_preserved_ module (C) Plot of the log_2_ fold change from the DMD muscle transcriptome (LFC_DMD_) and the BMP4-simulated C2C12 transcriptome (LFC_BMP4_) for genes present in the Turquoise_preserved_ module with an FDR < 0.05. Dashed lines indicate transcripts with LFC > 2. Selected transcripts are labelled demonstrating high LFC.

#### Module preservation analysis between DMD muscle and BMP4-stimulated muscle cells

We next used module preservation analysis to test whether the BMP4-stimulated muscle cell transcriptomic network was overlapping with the DMD muscle transcriptomic co-expression network. The Z-Summary statistic was used to quantify the degree to which the gene co-expression networks were preserved (**Table S17**). We found moderate module preservation of the Turquoise_preserved_ module consisting of 1,027 upregulated transcripts with a Z-Summary score of 4.7 (**Fig. 5B, Table S18**). The preserved genes with the highest kMEs in the human and C2C12 networks, respectively, were *SERPING1 and Aff3*. The transcripts with the highest LFC were *ADAMTS3, FMOD, ID4*, *PRELP*, *PTGFR SFRP2*, *SMAD8*, and *WHRN* (**Fig. 5C**). S*MAD8* was also the only *SMAD* in the preserved Turquoise_preserved_ module. No downregulated modules were significantly preserved. However, *UNC13C* [LFC_DMD_ = −6.39 ± 2.25, FDR = 3.67 x 10^-02^ LFC_BMP4_ = −2.99 ± 0.85, FDR = 2.0 x 10^-03^] and *ANO5* [LFC_DMD_ = −2.2 ± 0.4, FDR = 1.3 x 10^-05^, LFC_BMP4_ = −2.7 ± 0.3 FDR = 4.7 x 10^-15^] were the most downregulated transcripts in both transcriptomes.

#### qPCR of selected dysregulated targets

In BMP4-stimulated C2C12 muscle cells, we assessed several transcripts that were found to be highly dysregulated in the late-stage DMD muscle including *DHRS9*, *HCAR2*, and *UNC13C* (**Fig. 6A**). *Dhrs9* was downregulated by BMP4 stimulation in contrast to its upregulation in DMD muscle. *Hcar2*, which was the most upregulated late-stage DMD muscle transcript, was induced more than 2-fold following BMP4 stimulation of C2C12 muscle cells. *Unc13c*, one of the most downregulated transcripts in DMD muscle, was suppressed by more than 8-fold in BMP4-stimulated cells. The top 10 *Unc13c* connected transcripts in BMP4-stimulated muscle cells by WGCNA is shown (**Fig. 6B, Table S19**).

**Fig. 6.**
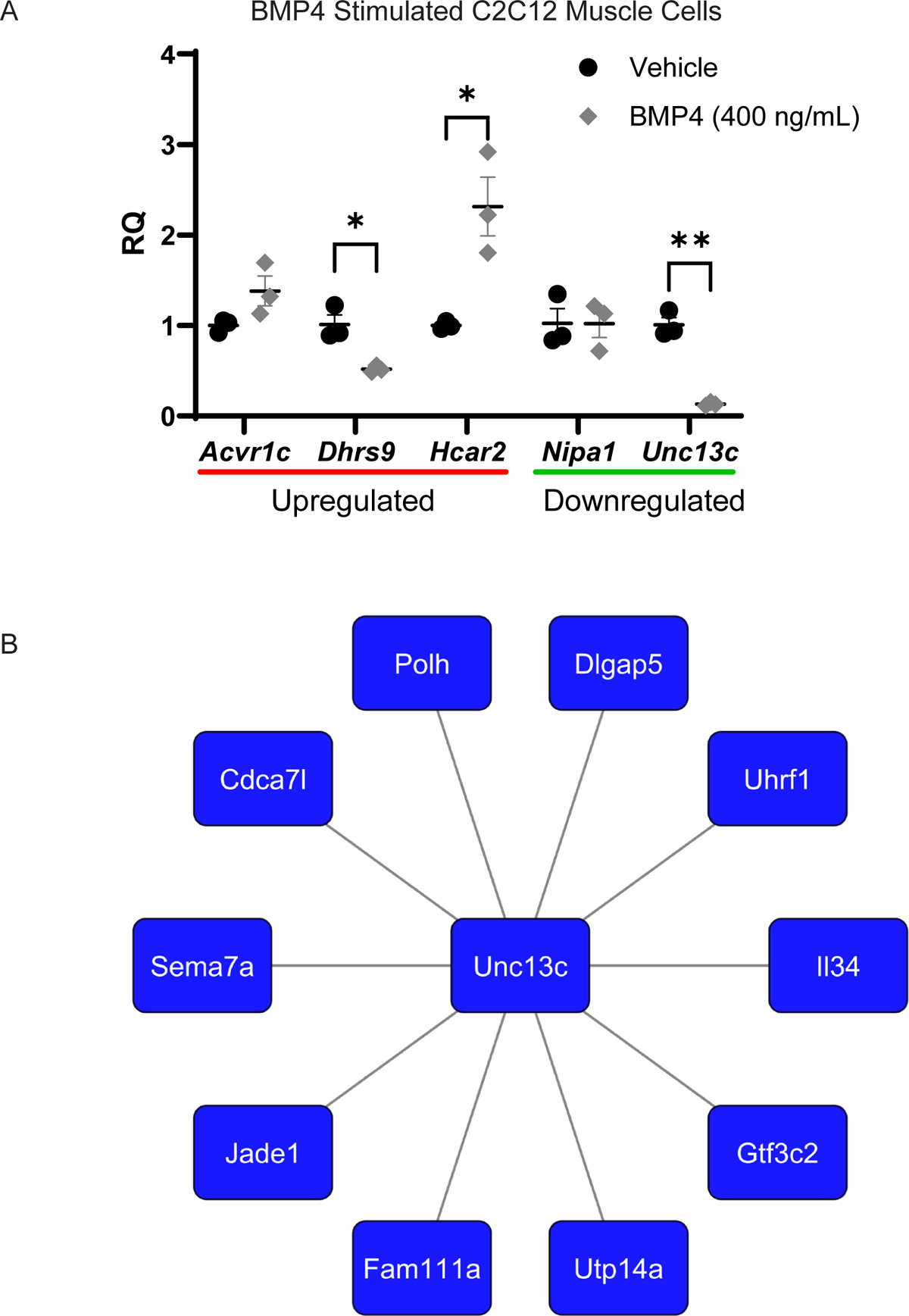
DMD dysregulated mRNA targets regulated by BMP4. (A) Plot shows mRNA expression levels (RQ) of *Acvr1c*, *Dhrs9*, *Hcar2, Nipa1,* and *Unc13c* after BMP4 stimulation of C2C12 muscle cell as measured by quantitative RT-PCR. Bars show mean ± SEM of 3 biological replicates per group. P-Values compared to vehicle * < 0.05 and ** < 0.01. RQ, relative quantity. Transcripts underlined in red and blue notes their upregulation or downregulation, respectively, in the DMD transcriptome. (B) *Unc13c* co-expressed genes with edge weight (from adjacency matrix) ≥ 50th percentile (0.387) of adjacency matrix values.

## Discussion

A deleterious role for BMP4 emerged from our prior studies, where we demonstrated that it induces and activates *SMAD8*, which then represses myomiRs: miR-1, miR-133a, and miR-133b ^8,15^. Based on these findings, we hypothesized that BMP4 signaling is part of a DMD disease-driving pathway. In support of this hypothesis, we found 1,027 differentially upregulated gene transcripts in the Turquoise_preserved_ module that were preserved between late-stage DMD skeletal muscle and BMP4-stimulated C2C12 muscle cells (**Fig. 5**, **Table S18**). This implies expression of these specific co-expressed transcripts in late-stage DMD skeletal muscle may be driven by BMP4 signaling. Within this preserved module, *SERPING1* and *Aff3* were identified as the top hub genes for the DMD and BMP4-induced transcriptomes, respectively. We also identify novel key genes with the highest fold-change in both transcriptomes*: ADAMTS3*, CPM, *FMOD, HCAR2, ID4*, *PTGFR*, *PRELP*, *SMAD8*, *SFRP2*, and *WHRN*. An increase in *SMAD8* was consistent with our prior study and was the only R-SMAD upregulated in either transcriptome. *SFRP2* was previously identified as highly upregulated in *mdx* and by muscle injury^29^. Further investigation of this preserved module will lay the foundation for future studies examining these pathways, their roles as drivers of DMD pathology, and the development of novel therapies to mitigate disease progression.

From our analysis, we identify three major processes relevant to DMD pathology: (1) immune dysfunction, (2) ECM remodeling, and (3) myogenic dysfunction. For the immune process, we find that TGFβ signaling continues to be an active pathway with upregulation of *TGFβ1*, *TGFβ3*, and the downstream signal transducer *SMAD8*^8^. The upregulation of TGFβ1 in BMP4-stimulated C2C12 muscle cells provides support for our hypothesis that this signaling pathway is operative in late-stage DMD muscle. Our previous finding that all three isoforms of TGF*β* induce and activate Smad8 raises the possibility of cross-activation of TGFβ1 downstream signaling via intracellular factors, potentially mediated through the R-Smads via non-canonical pathways^20,30,31^. This potential inter-connectivity has implications for therapeutic development given transcriptomic evidence of TGF signaling pathway redundancy, i.e. targeting of TGFβ1 or myostatin signaling alone may be insufficient to improve DMD disease trajectory^32,33^.

Additionally, *SERPING1* was identified as a hub gene in both the DMD and BMP4-stimulated muscle cell networks. *SERPING1* is involved in complement activation and encodes an inhibitor of complement activation (C1-INH)^35^. It plays an important role in suppressing inflammatory responses. It was previously identified as a hub gene in a microarray database of DMD muscle samples (GSE6011)^9^. The preservation of this hub gene in advanced stage DMD muscle suggests an ongoing role of complement activation in disease progression. We newly identify *HCAR2*, as the most upregulated transcript in the late-stage DMD muscle at 7.2-fold. HCAR2 is recognized as a repressor of inflammation and it is upregulated by TGFβ1^34,35^. In prior microarray studies from younger 5- to 9-year-old DMD males (GSE3307) *HCAR2* was minimally increased, suggesting it could serve as a novel biomarker for DMD progression and reflect an ongoing anti-inflammatory adaptative mechanism.

The anti-inflammatory effect of HCAR2 was previously demonstrated in several models including DMD primary myoblasts involving IL-6, in the *mdx* mouse, and macrophages via nicotinic acid via NFκB suppression^34–38^.

*Hcar2* was increased 2-fold by BMP4 stimulation of C2C12 muscle cells and WCGNA identified it as a part of Turquoise_BMP4_ module. This suggests close connectivity between BMP4 and HCAR2 and raises the possibility that BMP4 is an important regulator of HCAR2 in late-stage DMD muscle.

In contrast, we observed that *HLA-A* is the overall most downregulated transcript in late-stage DMD muscle. It is a major histocompatibility complex, class I, A molecule ubiquitously expressed and functioning in antigen presentation to immune system. A prior array study reports an upregulation of HLA genes, including *HLA-A*, in association with younger DMD biceps brachii muscles^39^. Thus, its downregulation in our samples suggests that attenuation of HLA genes, like *HLA-A*, are markers of disease progression. However, we did not find HLA genes to be reduced by BMP4 and this suggests other pathways drive this suppression. As further evidence of active pro-inflammatory and pro-fibrotic responses in late-stage DMD muscle, IPA identified activation of the Phagosome Formation CP and several upstream regulators; notably TGFβ1, IFNα/γ, and TNF. Taken together, immune-mediated mechanisms remain highly relevant in late-stage DMD, and the targeting of these members or their networks to promote anti-inflammatory effects may be an important therapeutic strategy even in advanced disease.

The second major process identified in late-stage DMD muscle was the ECM pathway which is linked to active remodeling, collagen deposition, and fibrofatty replacement of myofibers. The ADAM and ADAMTS families were highly upregulated. They function in ECM organization and collagen biosynthesis. *ADAMTS3, like ADAMTS2,* is a procollagen N-proteinases converting procollagens to collagen^40^. ADAM and ADAMTS upregulation promotes muscle fiber loss, fibrosis, and adipogenesis in *mdx* mice^41,42^. ADAM12, which is induced by TGFβ1, was the highest upregulated DMD muscle transcript. *ADAMTS3* was the highest induced transcript in BMP4-stimulated muscle cells. While WGCNA showed in DMD muscle that *ADAMTS2* had high connectivity to *SMAD8*. Taken together, these findings highlight a novel link between BMP4 signaling, SMAD8, and upregulation of ECM remodeling signaling via the ADAMTS family in late-stage DMD muscle.

MMPs are highly relevant modulators of the ECM in DMD and the *MMP9* transcript was amongst the highest upregulated at 3-fold. Through its degradation activity on the basement and sarcolemmal membranes, MMP-9 promotes myofiber injury, fibrosis, and impairs regeneration^43^. It also promotes TGFβ signaling by converting TGFβ to an active form via proteolytic processing. Prior array databases and other reports did not identify MMP9 as upregulated or active in younger DMD patient muscles (**Fig. 3B**)^44,45^. Although the prior array studies may not have included MMP9, our detection of increased MMP9 in older, advanced disease suggests that it is a marker of disease progression. MMP9 can be induced by IL-1β and TNFα / NFκB via SAF-1/AP-1 dependent mechanisms^46^. Indeed, serum MMP-9 has been identified as a potential biomarker of clinical progression with significantly higher levels detected in older DMD patients^47^. Interestingly, MMP9 knockout studies suggest a disease-mitigating role for MMP9 in the late dystrophic stage in contrast to earlier stages^48^. Our studies indicate that *MMP2*, *14*, *19*, and *23B* were commonly upregulated in DMD and by BMP4 and were in the Turquoise_preserved_ module. The overlap of BMP4-induced ADAMTS / MMP family members with the DMD transcriptome supports BMP4 as an important signaling pathway in the fibrotic conversion of late-stage DMD skeletal muscle. This interpretation is further supported by common IPA CPs that were activated which included ECM, Molecular Mechanisms of Cancer, and Phagosome Formation CPs.

Thirdly, we found significant evidence of pathways related to myogenesis and loss of muscle metabolic functions. While there was evidence of ongoing muscle regeneration with increased *MYOG*, *MYH3*, and *MYH8*, there was also downregulation of transcriptional regulators of myogenesis including *MEF2D* and *MYF6*. Downregulation of *DMD* and *CKM* transcripts was expected and consistent with DMD muscle pathology reflecting ongoing loss of myofibers. The top IPA identified canonical pathways and upstream regulators pertinent to muscle loss included activation of Mitochondrial Dysfunction and inhibition of metabolic categories (Electron Transport, Oxidative Phosphorylation, and Citric Acid). Prediction inhibition of the Mitochondrial Dysfunction CP was previously reported in younger DMD skeletal muscle microarrays (GSE3307); although with predicted Z-scores that were lower than our late-stage DMD suggesting worsening of mitochondrial dysfunction over age^14,27^. Overall, these patterns reflect loss of normal mitochondrial rich metabolic activity with concomitant activation of mitochondrial dysfunction in DMD (**Fig. 2A/B**).

Downregulated transcripts in the DMD muscle included three significant modules with the Blue_DMD_ module being the largest and containing the *DMD* transcript. Within the Blue_DMD_ module, we identified over 100 genes highly connected to DMD. *UNC13C* was one of the most downregulated transcripts in DMD muscle and this marked attenuation was strongly reproduced in BMP4-stimulated muscle cells indicating a major role for BMP4 signaling. *UNC13C* function is not well explored, but prior reports in *C. elegans* show *unc* mutations confer severe impairments in motility and aberrant neuromuscular junction morphometry^49,50^. *Unc13c* is also implicated in the promotion of myoblast differentiation and negatively regulated by TNFα^51^. Other co-downregulated transcripts in DMD*_blue_* module included *SGCB* and *MRPL39*. *SGCB* is associated with limb girdle muscular dystrophy and in DMD there is secondary loss of *SGCB* expression at the sarcolemma^28^. Notably, increasing *SGCB* expression is thought to improve dystrophic muscle health. While *MRPL39* is involved in protein synthesis vis-a-vis mitochondrial DNA maintenance and it is regulated by *TP53*^52,53^.

In addition to TGFβ1, several regulators were revealed with connections to BMP4 and TGF signaling. RICTOR, an assembly component of the MTOR complex, is increased by TGFβ1 and it interacts with BMP4^54–58^. Prior studies show that RICTOR increases activation of TP53^59,60^. *SERPING1*, which we and others identify as a key hub gene, is regulated by TP53^61^. Thus, these data suggest that TP53, TGβ1, BMP4, and SMAD8 may share signaling networks that are progressively upregulated in DMD. Additional study is required to understand how targeting of these common downstream effectors of the TGF superfamily impact DMD disease. For example, in the case of *SMAD8*, our work has shown it can have a deleterious impact in myogenesis and suggests that targeting it may be a more effective strategy for mitigating dysregulation of immune, ECM, and myogenic processes in DMD.

In summary, our study provides new insight into the transcriptome of late-stage DMD skeletal muscle. We found a significant overlap with the BMP4-induced transcriptome in muscle cells suggesting that BMP4 signaling plays a major role in shaping the molecular signature in this disease stage. Additionally, our study identified common and unique signatures in late-stage DMD compared to early stage with the emergence of several potential biomarkers of disease progression. Although a limitation of the study was small sample size, due to the scarcity of muscle samples available at this age, our findings provide a road map for future investigations into the role of BMP4 signaling and downstream factors like *HCAR2*, *SMAD8*, and *UNC13C* in promoting pro-fibrotic pathways (**Fig. 7**). Future investigation into Smad8 as a downstream transducer of BMP4 signaling may provide insight into the broader role of TGF signaling in promoting progression of DMD pathology.

**Fig. 7.**
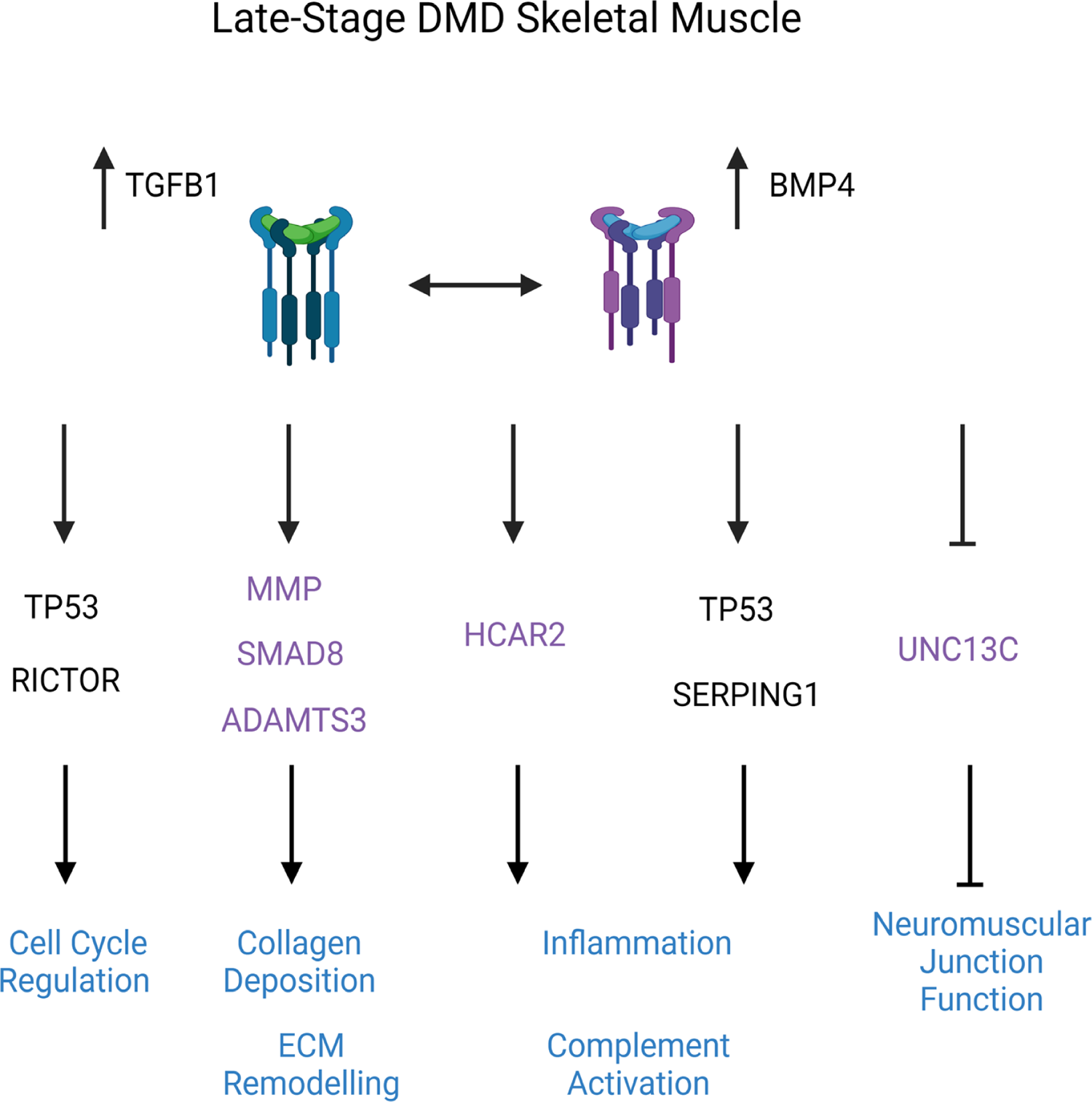
Model of late-stage DMD skeletal muscle transcriptomic changes in TGFβ signaling pathway. Blue text indicates disease processes or cellular function. Purple text indicates BMP4-induced regulation. Dashed arrows indicate activation by either pathway. Arrows indicate activation. Lines with blunt edges indicate inhibition.

## Acknowledgements

The authors would like to thank Michael Conklin, Scott Doyle, Shawn Gilbert, and Meagan Whatley for their invaluable technical assistance and collection of muscle samples. We also would like to thank the patients and families who generously participated in the study. This research was supported by grants from the National Institutes of Health (NIH) and other sponsors to authors as follows: H.S. is supported by the T32 UAB Center for Exercise Medicine. M.A.L. is supported by the National Institute of Neurological Disorders and Stroke (K08-NS120812) and UAB Center for Exercise Medicine. M.S.A. is supported by the National Institute of Child Health and Human Development (R01HD095897). P.H.K. is supported by NIH grants (R01NS092651, R21NS111275-01) and by the Department of Veterans Affairs (BX001148). M.N.M is supported by the National Heart, Lung, and Blood Institute (NHLBI) (R01 HL153460).

---

**Fig. 1S** Principal component analysis (PCA) of late-stage human DMD skeletal muscles and BMP4-stimulated C2C12 muscle cell transcriptomes. (A) PCA plot showing the separation between the DMD and non-DMD muscles along two dimensionless vectors PC1 and PC2. (B) PCA plot showing separation between the BMP4-stimulated C2C12 muscle cells compared with vehicle (VEH) along two dimensionless vectors PC1 and PC2. Each group consists of 3 biological replicates (N=3).

**Fig. 2S** Sensitivity analysis of BMP4-stimulated C2C12 myoblast transcriptomes with and without the outlier. (A) Plot shows the log_2_ fold change (LFC) comparing with (outlier in) and without outlier (outlier out). (B) Plot shows the adjusted (Padj) P-Value comparing with outlier in and outlier out.

## List of Supplementary Tables

**Table S0A** All expressed genes DMD vs. non-DMD skeletal muscles with normalized counts.

**Table S0B** All expressed genes BMP4-stimulated C2C12 muscle cells vs. vehicle with normalized counts.

**Table S1** Differential gene expression analysis of DMD skeletal muscles (FDR < 0.05).

**Table S2** Canonical Pathways in DMD skeletal muscle transcriptome using Ingenuity Pathway Analysis.

**Table S3** Upstream regulators in DMD skeletal muscle transcriptome using Ingenuity Pathway Analysis.

**Table S4** Analysis Match of DMD skeletal muscle transcriptome using Ingenuity Pathway Analysis.

**Table S5** Modules identified in the DMD skeletal muscle transcriptome using weighted gene co-expression network analysis.

**Table S6** Genes in significant upregulated modules from the weighted gene co-expression network analysis of the Duchenne muscular dystrophy skeletal muscle transcriptome

**Table S7** Genes in significant down regulated modules from the weighted gene co-expression network analysis of the Duchenne muscular dystrophy skeletal muscle transcriptome

**Table S8** Genes highly co-expressed with SMAD8 (based on adjacency matrix cutoff of ≥ 0.40) neighbors in turquoise module in Duchenne muscular dystrophy skeletal muscle transcriptome.

**Table S09** Genes highly co-expressed with HCAR2 (based on adjacency matrix cutoff of ≥0.40) neighbors in turquoise module in Duchenne muscular dystrophy skeletal muscle transcriptome using weighted gene co-expression network analysis.

**Table S10** Genes highly co-expressed with DMD (based on adjacency matrix cutoff of ≥ 0.45) from the blue module in the Duchenne muscular dystrophy skeletal muscle transcriptome.

**Table S11** Differential gene expression analysis of BMP4-stimulated C2C12 myoblasts transcriptome using DESeq method (FDR < 0.05).

**Table S12** Canonical Pathways in BMP4-stimulated C2C12 myoblasts transcriptome using Ingenuity Pathway Analysis.

**Table S13** Upstream regulators in BMP4-stimulated C2C12 myoblasts transcriptome using Ingenuity Pathway Analysis.

**Table S14** Modules identified in the BMP4-stimulated C2C12 myoblasts transcriptome using weighted gene co-expression network analysis.

**Table S15** Genes in the turquoise module of the BMP4-stimulated C2C12 muscle cells transcriptome using weighted gene co-expression network analysis.

**Table S16** Genes in the blue module of the BMP4-stimulated C2C12 muscle cells transcriptome using weighted gene co-expression network analysis.

**Table S17** Module preservation analysis between DMD muscle and BMP4-stimulated myoblasts transcriptomes.

**Table S18** Fold-change of preserved genes in Turquoise modules of DMD muscle and BMP4-transcriptomes.

**Table S19** Genes highly co-expressed with *Unc13c* (based on top 50% of weights in adjacency matrix) in blue module in BMP4-stimulated C2C12 muscle cells transcriptome using weighted gene co-expression network analysis.

## References

1. Hoffman EP, Brown RH, Jr., Kunkel LM. Dystrophin: the protein product of the Duchenne muscular dystrophy locus. Cell. Dec 24 1987;51(6):919–28. doi:10.1016/0092-8674(87)90579-4

2. Rosenberg AS, Puig M, Nagaraju K, et al. Immune-mediated pathology in Duchenne muscular dystrophy. Sci Transl Med. Aug 5 2015;7(299):299rv4. doi:10.1126/scitranslmed.aaa7322

3. Broomfield J, Hill M, Guglieri M, Crowther M, Abrams K. Life Expectancy in Duchenne Muscular Dystrophy: Reproduced Individual Patient Data Meta-analysis. Neurology. Dec 7 2021;97(23):e2304–e2314. doi:10.1212/wnl.0000000000012910

4. Nishio H, Takeshima Y, Narita N, et al. Identification of a novel first exon in the human dystrophin gene and of a new promoter located more than 500 kb upstream of the nearest known promoter. J Clin Invest. Sep 1994;94(3):1037–42. doi:10.1172/jci117417

5. Mázala DA, Novak JS, Hogarth MW, et al. TGF-β-driven muscle degeneration and failed regeneration underlie disease onset in a DMD mouse model. JCI Insight. Mar 26 2020;5(6)doi:10.1172/jci.insight.135703

6. Rosenberg AS, Puig M, Nagaraju K, et al. Immune-mediated pathology in Duchenne muscular dystrophy. Science translational medicine. 07/15/nihms-submitted 05/14/pmc-release 2015;7(299):299rv4–299r<otherinfo>v4. doi:10.1126/scitranslmed.aaa7322</otherinfo>

7. Chen YW, Nagaraju K, Bakay M, et al. Early onset of inflammation and later involvement of TGFbeta in Duchenne muscular dystrophy. Neurology. Sep 27 2005;65(6):826–34. doi:10.1212/01.wnl.0000173836.09176.c4

8. Lopez MA, Si Y, Hu XZ, et al. Smad8 Is Increased in Duchenne Muscular Dystrophy and Suppresses miR-1, miR-133a, and miR-133b. International Journal of Molecular Sciences. Jul 2022;23(14):7515. doi:ARTN 7515 10.3390/ijms23147515

9. Wei Y, Su Q, Li X. Identification of hub genes related to Duchenne muscular dystrophy by weighted gene co-expression network analysis. Medicine (Baltimore). Dec 30 2022;101(52):e32603. doi:10.1097/md.0000000000032603

10. Chen Y-W, Zhao P, Borup R, Hoffman EP. Expression Profiling in the Muscular Dystrophies: Identification of Novel Aspects of Molecular Pathophysiology. Journal of Cell Biology. 2000;151(6):1321–1336. doi:10.1083/jcb.151.6.1321

11. Bakay M, Zhao P, Chen J, Hoffman EP. A web-accessible complete transcriptome of normal human and DMD muscle. Neuromuscular Disorders. 2002/10/01/ 2002;12:S125–S141. 10.1016/S0960-8966(02)00093-7

12. Haslett JN, Sanoudou D, Kho AT, et al. Gene expression comparison of biopsies from Duchenne muscular dystrophy (DMD) and normal skeletal muscle. Proceedings of the National Academy of Sciences. 2002;99(23):15000–15005. doi:10.1073/pnas.192571199

13. Pescatori M, Broccolini A, Minetti C, et al. Gene expression profiling in the early phases of DMD: a constant molecular signature characterizes DMD muscle from early postnatal life throughout disease progression. Faseb j. Apr 2007;21(4):1210–26. doi:10.1096/fj.06-7285com

14. Dadgar S, Wang Z, Johnston H, et al. Asynchronous remodeling is a driver of failed regeneration in Duchenne muscular dystrophy. J Cell Biol. Oct 13 2014;207(1):139–58. doi:10.1083/jcb.201402079

15. Si Y, Cui X, Kim S, et al. Smads as muscle biomarkers in amyotrophic lateral sclerosis. Ann Clin Transl Neurol. Oct 2014;1(10):778–87. doi:10.1002/acn3.117

16. Dobin A, Davis CA, Schlesinger F, et al. STAR: ultrafast universal RNA-seq aligner. Bioinformatics. Jan 1 2013;29(1):15–21. doi:10.1093/bioinformatics/bts635

17. Coenen-Stass AML, Sork H, Gatto S, et al. Comprehensive RNA-Sequencing Analysis in Serum and Muscle Reveals Novel Small RNA Signatures with Biomarker Potential for DMD. Mol Ther Nucleic Acids. Dec 7 2018;13:1–15. doi:10.1016/j.omtn.2018.08.005

18. Love MI, Huber W, Anders S. Moderated estimation of fold change and dispersion for RNA-seq data with DESeq2. Genome Biol. 2014;15(12):550. doi:10.1186/s13059-014-0550-8

19. Matsye P, Zheng L, Si Y, et al. HuR promotes the molecular signature and phenotype of activated microglia: Implications for amyotrophic lateral sclerosis and other neurodegenerative diseases. Glia. Jun 2017;65(6):945–963. doi:10.1002/glia.23137

20. Si Y, Cui X, Crossman DK, et al. Muscle microRNA signatures as biomarkers of disease progression in amyotrophic lateral sclerosis. Neurobiol Dis. Jun 2018;114:85–94. doi:10.1016/j.nbd.2018.02.009

21. Kramer A, Green J, Pollard J, Jr., Tugendreich S. Causal analysis approaches in Ingenuity Pathway Analysis. Bioinformatics. Feb 15 2014;30(4):523–30. doi:10.1093/bioinformatics/btt703

22. Zhang B, Horvath S. A general framework for weighted gene co-expression network analysis. Stat Appl Genet Mol Biol. 2005;4:Article17. doi:10.2202/1544-6115.1128

23. Langfelder P, Horvath S. WGCNA: an R package for weighted correlation network analysis. BMC Bioinformatics. Dec 29 2008;9(1):559. doi:10.1186/1471-2105-9-559

24. Langfelder P, Luo R, Oldham MC, Horvath S. Is my network module preserved and reproducible? PLoS Comput Biol. Jan 20 2011;7(1):e1001057. doi:10.1371/journal.pcbi.1001057

25. Wise A, Foord SM, Fraser NJ, et al. Molecular Identification of High and Low Affinity Receptors for Nicotinic Acid*. Journal of Biological Chemistry. 2003/03/14/ 2003;278(11):9869–9874. 10.1074/jbc.M210695200

26. Ehrlich KC, Lacey M, Ehrlich M. Epigenetics of Skeletal Muscle-Associated Genes in the ASB, LRRC, TMEM, and OSBPL Gene Families. Epigenomes. Jan 30 2020;4(1)doi:10.3390/epigenomes4010001

27. Bakay M, Wang Z, Melcon G, et al. Nuclear envelope dystrophies show a transcriptional fingerprint suggesting disruption of Rb-MyoD pathways in muscle regeneration. Brain. Apr 2006;129(Pt 4):996–1013. doi:10.1093/brain/awl023

28. Durbeej M, Cohn RD, Hrstka RF, et al. Disruption of the beta-sarcoglycan gene reveals pathogenetic complexity of limb-girdle muscular dystrophy type 2E. Mol Cell. Jan 2000;5(1):141–51. doi:10.1016/s1097-2765(00)80410-4

29. Hara T, Nakayama Y, Nara N. [Regenerative medicine of skeletal muscle]. Rinsho Shinkeigaku. Nov 2005;45(11):880–2.

30. Radhakrishnan K, Luu M, Iaria J, et al. Activin and BMP Signalling in Human Testicular Cancer Cell Lines, and a Role for the Nucleocytoplasmic Transport Protein Importin-5 in Their Crosstalk. Cells. Mar 24 2023;12(7)doi:10.3390/cells12071000

31. Si Y, Kim S, Cui X, et al. Transforming Growth Factor Beta (TGF-beta) Is a Muscle Biomarker of Disease Progression in ALS and Correlates with Smad Expression. PLoS One. 2015;10(9):e0138425. doi:10.1371/journal.pone.0138425

32. Campbell C, McMillan HJ, Mah JK, et al. Myostatin inhibitor ACE-031 treatment of ambulatory boys with Duchenne muscular dystrophy: Results of a randomized, placebo-controlled clinical trial. Muscle & nerve. 2017;55(4):458–464. 10.1002/mus.25268

33. Guiraud S, Aartsma-Rus A, Vieira NM, Davies KE, van Ommen GJ, Kunkel LM. The Pathogenesis and Therapy of Muscular Dystrophies. Annu Rev Genomics Hum Genet. 2015;16:281–308. doi:10.1146/annurev-genom-090314-025003

34. Hamby ME, Coppola G, Ao Y, Geschwind DH, Khakh BS, Sofroniew MV. Inflammatory mediators alter the astrocyte transcriptome and calcium signaling elicited by multiple G-protein-coupled receptors. J Neurosci. Oct 17 2012;32(42):14489–510. doi:10.1523/jneurosci.1256-12.2012

35. Kalkan H, Pagano E, Paris D, et al. Targeting gut dysbiosis against inflammation and impaired autophagy in Duchenne muscular dystrophy. EMBO Mol Med. Mar 8 2023;15(3):e16225. doi:10.15252/emmm.202216225

36. Li G, Deng X, Wu C, et al. Distinct Kinetic and Spatial Patterns of Protein Kinase C (PKC)- and Epidermal Growth Factor Receptor (EGFR)-dependent Activation of Extracellular Signal-regulated Kinases 1 and 2 by Human Nicotinic Acid Receptor GPR109A*. Journal of Biological Chemistry. 2011/09/09/ 2011;286(36):31199–31212. 10.1074/jbc.M111.241372

37. Zandi-Nejad K, Takakura A, Jurewicz M, et al. The role of HCA2 (GPR109A) in regulating macrophage function. Faseb j. Nov 2013;27(11):4366–74. doi:10.1096/fj.12-223933

38. Li G, Deng X, Wu C, et al. Distinct kinetic and spatial patterns of protein kinase C (PKC)- and epidermal growth factor receptor (EGFR)-dependent activation of extracellular signal-regulated kinases 1 and 2 by human nicotinic acid receptor GPR109A. J Biol Chem. Sep 9 2011;286(36):31199–212. doi:10.1074/jbc.M111.241372

39. Noguchi S, Tsukahara T, Fujita M, et al. cDNA microarray analysis of individual Duchenne muscular dystrophy patients. Hum Mol Genet. Mar 15 2003;12(6):595–600.

40. Porter S, Clark IM, Kevorkian L, Edwards DR. The ADAMTS metalloproteinases. Biochem J. Feb 15 2005;386(Pt 1):15–27. doi:10.1042/bj20040424

41. Jørgensen LH, Jensen CH, Wewer UM, Schrøder HD. Transgenic overexpression of ADAM12 suppresses muscle regeneration and aggravates dystrophy in aged mdx mice. Am J Pathol. Nov 2007;171(5):1599–607. doi:10.2353/ajpath.2007.070435

42. Solomon E, Li H, Duhachek Muggy S, Syta E, Zolkiewska A. The role of SnoN in transforming growth factor beta1-induced expression of metalloprotease-disintegrin ADAM12. J Biol Chem. Jul 16 2010;285(29):21969–77. doi:10.1074/jbc.M110.133314

43. Ogura Y, Tajrishi MM, Sato S, Hindi SM, Kumar A. Therapeutic potential of matrix metalloproteinases in Duchenne muscular dystrophy. Front Cell Dev Biol. 2014;2:11. doi:10.3389/fcell.2014.00011

44. von Moers A, Zwirner A, Reinhold A, et al. Increased mRNA expression of tissue inhibitors of metalloproteinase-1 and −2 in Duchenne muscular dystrophy. Acta Neuropathol. Mar 2005;109(3):285–93. doi:10.1007/s00401-004-0941-0

45. Zanotti S, Saredi S, Ruggieri A, et al. Altered extracellular matrix transcript expression and protein modulation in primary Duchenne muscular dystrophy myotubes. Matrix Biol. Oct 2007;26(8):615–24. doi:10.1016/j.matbio.2007.06.004

46. Ray A, Bal BS, Ray BK. Transcriptional induction of matrix metalloproteinase-9 in the chondrocyte and synoviocyte cells is regulated via a novel mechanism: evidence for functional cooperation between serum amyloid A-activating factor-1 and AP-1. J Immunol. Sep 15 2005;175(6):4039–48. doi:10.4049/jimmunol.175.6.4039

47. Nadarajah VD, van Putten M, Chaouch A, et al. Serum matrix metalloproteinase-9 (MMP-9) as a biomarker for monitoring disease progression in Duchenne muscular dystrophy (DMD). Neuromuscul Disord. Aug 2011;21(8):569–78. doi:10.1016/j.nmd.2011.05.011

48. Shiba N, Miyazaki D, Yoshizawa T, et al. Differential roles of MMP-9 in early and late stages of dystrophic muscles in a mouse model of Duchenne muscular dystrophy. Biochim Biophys Acta. Oct 2015;1852(10 Pt A):2170–82. doi:10.1016/j.bbadis.2015.07.008

49. Brose N, Hofmann K, Hata Y, Sudhof TC. Mammalian homologues of Caenorhabditis elegans unc-13 gene define novel family of C2-domain proteins. J Biol Chem. Oct 20 1995;270(42):25273–80. doi:10.1074/jbc.270.42.25273

50. Varoqueaux F, Sons MS, Plomp JJ, Brose N. Aberrant morphology and residual transmitter release at the Munc13-deficient mouse neuromuscular synapse. Mol Cell Biol. Jul 2005;25(14):5973–84. doi:10.1128/MCB.25.14.5973-5984.2005

51. Meyer SU, Krebs S, Thirion C, Blum H, Krause S, Pfaffl MW. Tumor Necrosis Factor Alpha and Insulin-Like Growth Factor 1 Induced Modifications of the Gene Expression Kinetics of Differentiating Skeletal Muscle Cells. PLoS One. 2015;10(10):e0139520. doi:10.1371/journal.pone.0139520

52. Mak TW, Hauck L, Grothe D, Billia F. p53 regulates the cardiac transcriptome. Proc Natl Acad Sci U S A. Feb 28 2017;114(9):2331–2336. doi:10.1073/pnas.1621436114

53. O’Brien TW, Fiesler SE, Denslow ND, et al. Mammalian mitochondrial ribosomal proteins (2). Amino acid sequencing, characterization, and identification of corresponding gene sequences. J Biol Chem. Dec 17 1999;274(51):36043–51. doi:10.1074/jbc.274.51.36043

54. Lamming DW, Demirkan G, Boylan JM, et al. Hepatic signaling by the mechanistic target of rapamycin complex 2 (mTORC2). Faseb j. Jan 2014;28(1):300–15. doi:10.1096/fj.13-237743

55. Rosenbluh J, Mercer J, Shrestha Y, et al. Genetic and Proteomic Interrogation of Lower Confidence Candidate Genes Reveals Signaling Networks in β-Catenin-Active Cancers. Cell Syst. Sep 28 2016;3(3):302–316.e4. doi:10.1016/j.cels.2016.09.001

56. Serrano I, McDonald PC, Lock FE, Dedhar S. Role of the integrin-linked kinase (ILK)/Rictor complex in TGFβ-1-induced epithelial-mesenchymal transition (EMT). Oncogene. Jan 3 2013;32(1):50–60. doi:10.1038/onc.2012.30

57. Samarakoon R, Overstreet JM, Higgins PJ. TGF-β signaling in tissue fibrosis: redox controls, target genes and therapeutic opportunities. Cell Signal. Jan 2013;25(1):264–8. doi:10.1016/j.cellsig.2012.10.003

58. Zhu D, Wu J, Spee C, Ryan SJ, Hinton DR. BMP4 mediates oxidative stress-induced retinal pigment epithelial cell senescence and is overexpressed in age-related macular degeneration. J Biol Chem. Apr 3 2009;284(14):9529–39. doi:10.1074/jbc.M809393200

59. Jung SH, Hwang HJ, Kang D, et al. mTOR kinase leads to PTEN-loss-induced cellular senescence by phosphorylating p53. Oncogene. Mar 2019;38(10):1639–1650. doi:10.1038/s41388-018-0521-8

60. Julien LA, Carriere A, Moreau J, Roux PP. mTORC1-activated S6K1 phosphorylates Rictor on threonine 1135 and regulates mTORC2 signaling. Mol Cell Biol. Feb 2010;30(4):908–21. doi:10.1128/mcb.00601-09

61. Wang L, Wu Q, Qiu P, et al. Analyses of p53 target genes in the human genome by bioinformatic and microarray approaches. J Biol Chem. Nov 23 2001;276(47):43604–10. doi:10.1074/jbc.M106570200

