## Supplementary figures and images for "Late-Stage Skeletal Muscle Transcriptome in Duchenne muscular dystrophy shows a BMP4-Induced Molecular Signature"

### Supplemental Figures

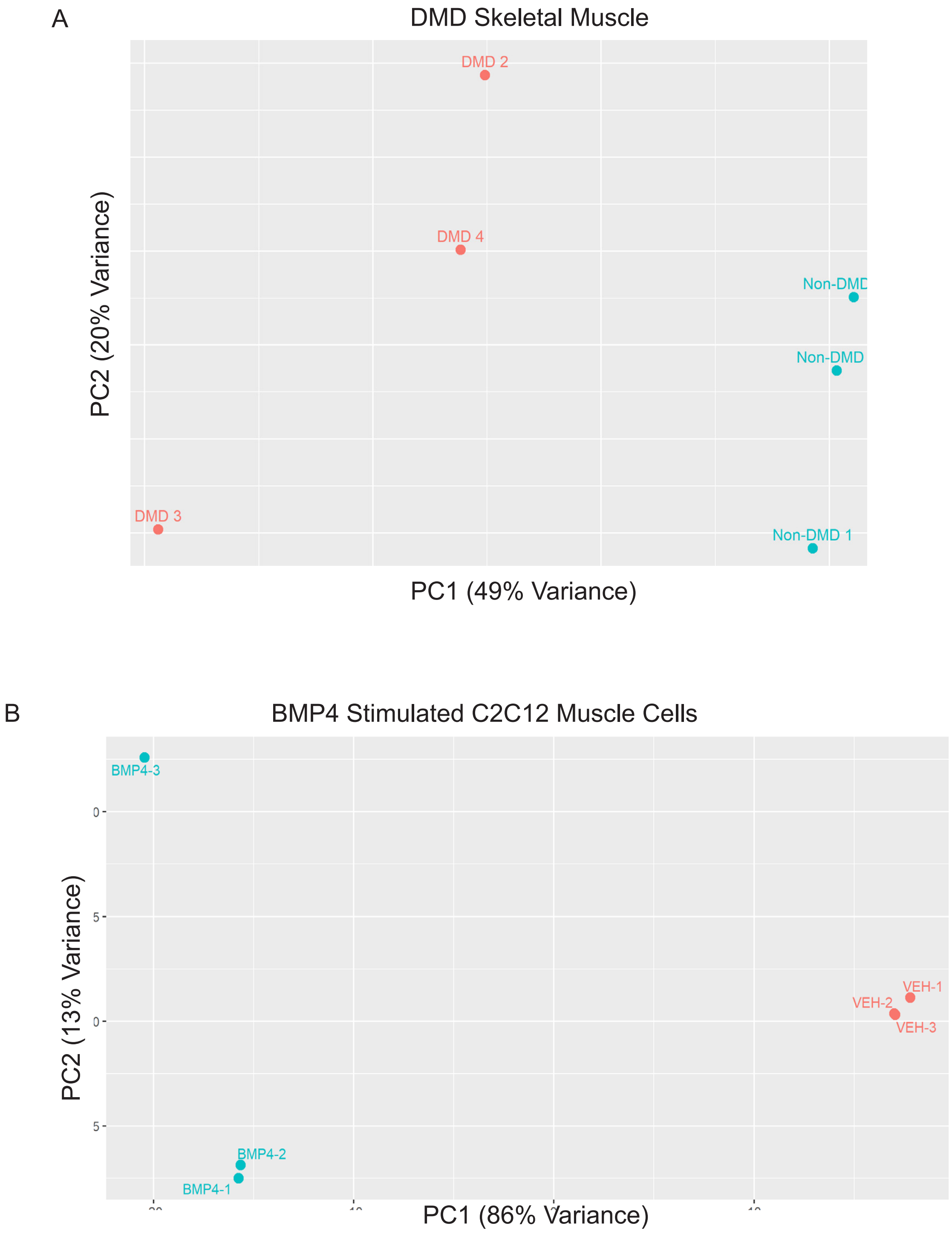

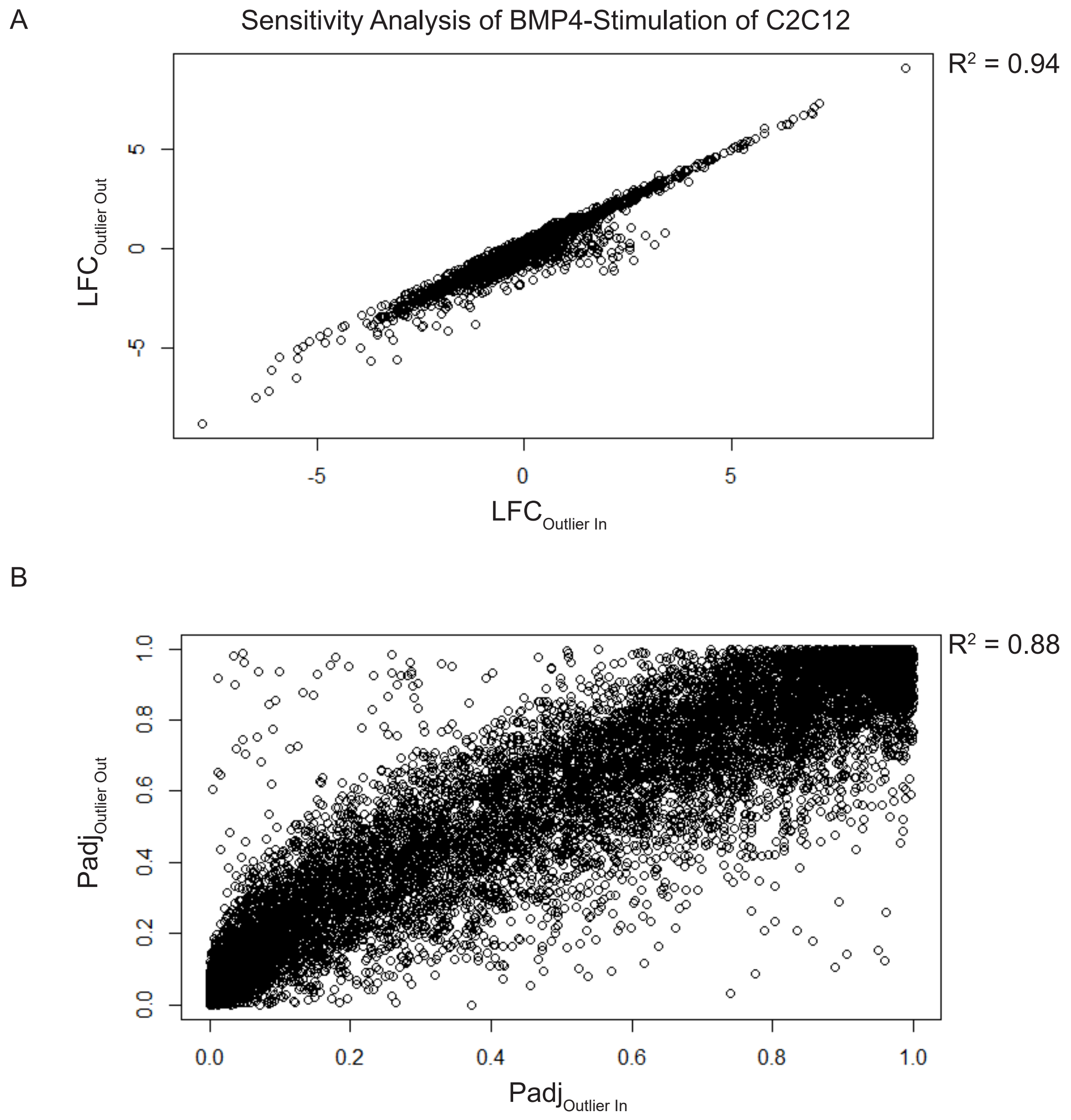
